# Disrupted bone microenvironment and immune recovery following total body irradiation in a murine model

**DOI:** 10.1101/2025.03.17.643638

**Authors:** Tibor Sághy, Priti Gupta, Karin Horkeby, Petra Henning, Claes Ohlsson, Carmen Corciulo, Marie K Lagerquist, Andrei S. Chagin, Cecilia Engdahl

**Author notes:** Corresponding author Cecilia Engdahl PhD, Department of Rheumatology and Inflammation Research Institute of Medicine, University of Gothenburg Gothenburg, Sweden.

## Abstract

Irradiation is an effective therapy for killing cancer cells and a critical preparative regimen for hematopoietic stem cell transplantation (HSCT). However, irradiation causes side effects on healthy tissue, disrupting bone tissue and bone marrow homeostasis and leading to skeletal and immune dysfunction. To investigate these effects, we utilized a female murine model to examine the mechanisms underlying skeletal damage caused by total body irradiation. Experiments were conducted over a 12-week period of total body irradiation and HSCT, complemented by an acute study involving only total body irradiation.

Irradiation led to a transient depletion of bone marrow immune cells, followed by a late increased bone marrow cellularity 12 weeks after irradiation and HSCT compared to naive, indicative of late-stage local immune induction. Cortical and trabecular bone damage emerged two weeks post-irradiation and HSCT and persisted throughout the study. This bone damage was accompanied by a sustained increase in bone marrow adiposity. Accumulation of apoptotic cells was observed in the bone marrow within six hours post-irradiation and persisted up to 12 weeks post-irradiation and HSCT. Accompanied by elevated local expression of the pro-apoptotic BAX gene and a gene elevated with the clearance of apoptotic cells, TGF-β1. *In vitro* studies revealed that macrophages and pre-osteoclasts, but not fully differentiated osteoclasts, efficiently cleared apoptotic cells with an elevation of TGF-β1 in culture supernatant. While the clearance of apoptotic cells and associated TGF-β1 signaling were evident, their direct role in skeletal outcomes remains unclear.

These findings suggest that persistent apoptotic cells contribute to impaired bone remodeling, possibly influencing osteoclast function and the bone microenvironment. Further research is warranted to explore whether targeting apoptotic cell clearance could reduce bone damage and support skeletal recovery following irradiation.

## Introduction

Irradiation therapy remains a fundamental component in the management of various cancers, with more than half of all cancer patients receiving radiotherapy, including total body irradiation (TBI)^1^. Despite its effectiveness in eradicating hematopoietic cells before hematopoietic stem cell transplantation (HSCT) and reducing tumor size, there are concerns about affecting healthy surrounding tissues, often leading to immediate and long-term adverse effects^2^. Skeletal tissue is particularly vulnerable to radiosensitivity due to the high calcium content^3,4^. Irradiation can reduce bone mineral density and disrupt bone microarchitecture^5–7^, but the timeline and underlying mechanisms are unclear. This weakened bone tissue significantly increases the risk of fractures^7,8^, contributing to higher morbidity and mortality among cancer patients and posing a substantial challenge to overall patient outcomes^9^.

Bone marrow hematopoietic stem cells (HSCs), essential for regenerating mature blood cells, are highly susceptible to irradiation. Even low doses of TBI can drastically lower lymphocyte counts, while high doses destroy both mature lymphocytes and HSCs, and HSCT is necessary for long-term survival^10^. Irradiation triggers extensive cellular damage through oxidative stress and DNA damage, leading to massive apoptosis. The clearance of apoptotic cells, i.e., efferocytosis, is primarily performed by professional phagocytes, such as macrophages and dendritic cells^11,12,13^. The maturation of these cells is compromised after irradiation, possibly influencing the efferocytosis capacity.

Osteoclasts originate from monocytes^14^. Monocytes exhibit a relative radiation resistance, but like other immune cells, they temporarily reduce their numbers after radiation^15^. Since monocytes serve as the precursor cells for osteoclasts, the reduction limits osteoclast formation and temporarily impairs bone resorption. However, as monocyte populations recover directly, the increase can lead to a surge in bone resorption in the later stages of recovery from irradiation^16^. Like other monocytes, osteoclasts can phagocytose pathogens and act as antigen-presenting cells in bone marrow^17,18^. Apoptotic osteoblast and bone lining could stimulate osteoclasts during normal bone remodeling with microfractures^19,20^ ^21,22^. However, the mechanisms of how apoptotic cells regulate the osteoclast activity and if a massive induction of apoptotic cells would influence the osteoclast remains unclear.

Moreover, radiation exposure initiates an inflammatory response within the bone microenvironment^23^. Inflammation initiates a local induction of pro-inflammatory cytokines^24^ and transforming growth factor-beta (TGF-β1)^25,26^. These mediators can affect bone hemostasis and regulate osteoclast activity and differentiation^14,27,28^.

Radiation-induced damage also triggers the deregulation of mesenchymal stem cells (MSC). It shifts MSC differentiation toward adipogenesis and leads to increased marrow adiposity, thereby reducing the capability to form osteoblasts, which collectively impair bone regeneration^29,30^.

We hypothesize that apoptotic cells regulate the bone microenvironment and modulate bone homeostasis, particularly after total body irradiation. Therefore, our study aims to characterize changes in immune cells, skeletal tissues, and the role of apoptotic cells in osteoclastogenesis in a mouse model. By elucidating the interactions, we seek to uncover mechanisms underlying the disruption of bone homeostasis post-irradiation.

## Material and methods

### Mice

8 weeks old female C57BL/6 wild-type mice were obtained from Janvier Labs (Le Genest Saint Isle, France) and maintained at the Sahlgrenska Academy, Laboratory of Experimental Biomedicine, Gothenburg. The mice were housed in standard cages, kept at 22°C, and a 12-hour light-dark cycle. They had free access to Teklad Diet 2016 (Envigo, Indianapolis, IN, USA) and tap water. Starting one week before irradiation and until two weeks post-irradiation, both control and irradiation mice received Enrofloxacin (0.6 mg/ml, Elanco Denmark ApS) in their drinking water as an antibiotic treatment. Before euthanasia, the final body mass of the mice was recorded. Mice were anesthetized with a Ketador/Dexdomitor vet (Richter Pharma/Orion Pharma), bled from the axillary vein, and euthanasia was performed via cervical dislocation. Organ weights were determined and normalized to body weight for analysis. Tissue samples were snap-frozen in liquid nitrogen and stored at −80°C, while bones were fixed in 4% paraformaldehyde (PFA) for 48 hours, followed by 70% ethanol and then stored at room temperature until subsequent analysis. The animal procedures complied with the guidelines of the Animal Ethics Committee of Gothenburg (3230–2020).

### Irradiation and hematopoietic stem cell transplantation (HSCT)

For the long-term irradiation experiments followed by HCST performed the same day (detailed visual irradiation scheme is in Fig. 1a), a 9 Gy lethal dose of total body X-ray irradiation was administered to the mice in two fractions of 4.5 Gy dose with a 4-hour interval between each fraction. For the 48-hour acute irradiation experiment, a 9 Gy lethal dose of total body X-ray irradiation was administered in one fraction (a detailed visual irradiation scheme is in Fig. 4a). Irradiation was performed using an RS2000 irradiator with a 0.3 mm copper filter and X-ray tube settings of 160 kV and 25 mA (1.59 Gy/min) (Rad Source Technologies, Buford, GA, USA).

**Fig. 1:**
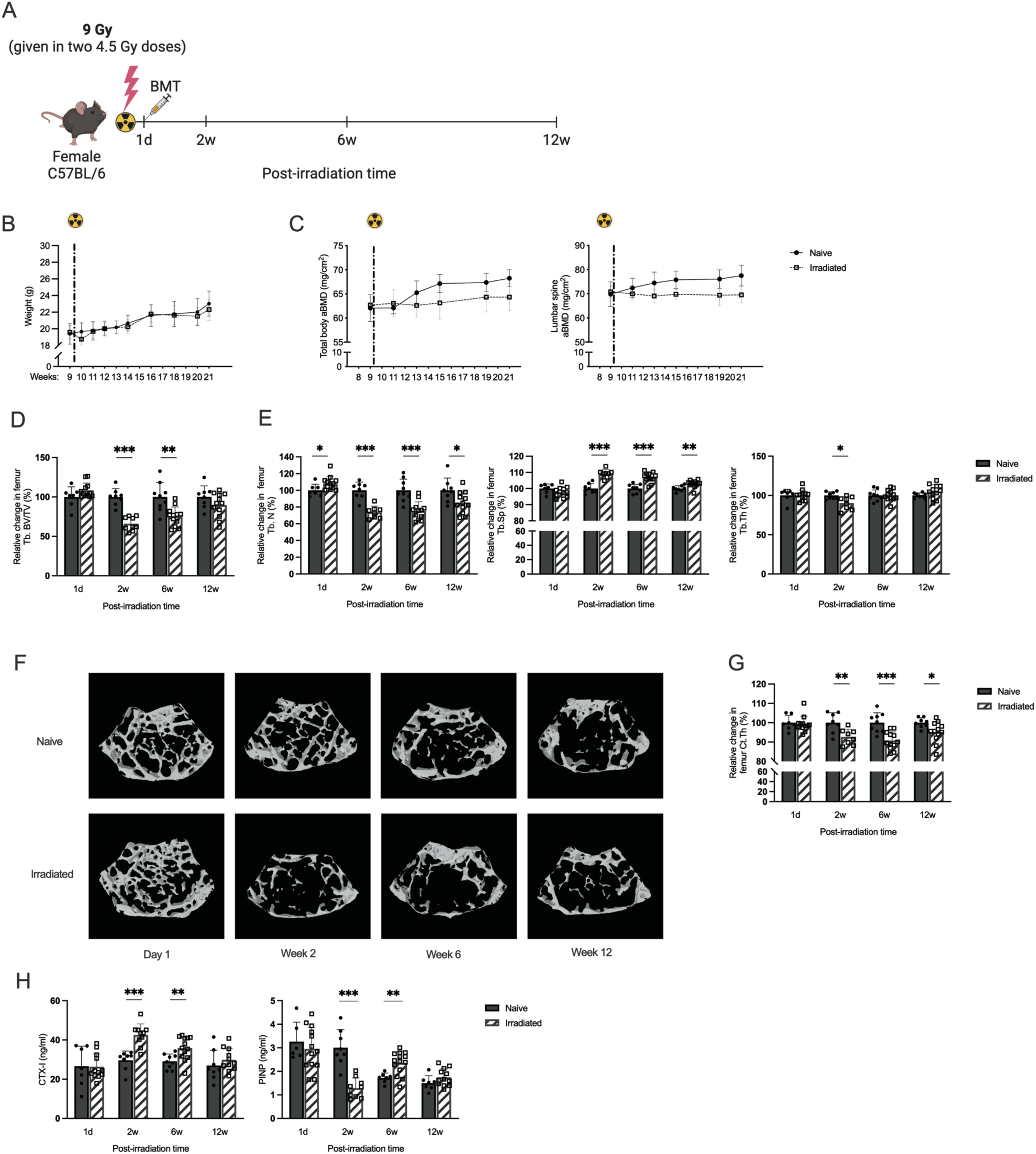
Longitudinal analysis of changes in bone parameters microstructure in response to irradiation and hematopoietic stem cell transplantation (HSCT). **A**. Schematic diagram showing experiment design. **B**. Changes in body weight over the twelve-week experiment. A significant difference was observed with Students t-test one week after irradiation and HSCT, age 10 weeks. The dotted line indicates the time of irradiation and HSCT. **C**. Areal bone mineral density (aBMD), total body, and lumbar spine vertebrae (LS) 3–6 in irradiated and control mice. A significant difference was observed with the Students’ t-test 4-weeks after irradiation and HSCT and persisted through the whole 12 weeks period, mice aged 13-21 weeks **D**. Micro-computed tomography (μCT) analysis of trabecular bone volume per total volume (Tb. BV/TV) in the femur. **E**. Trabecular number (Tb.N), trabecular separation (Tb.Sp), and trabecular thickness (Tb.Th) in the femur. **F**. Representative images of trabecular bone in femur. **G**. μCT measurements of femoral cortical thickness. **H**. Serum levels of C-terminal telopeptide (CTX, a bone resorption marker) and procollagen type I N-terminal propeptide (PINP, a bone formation marker). Statistical differences between irradiated and HSCT compared to naive mice at each time point were evaluated using Student’s t-test. Sample sizes ranged from n=8-12. Data are presented as mean ± SD. Significance levels are indicated as *P < 0.05, **P < 0.01, ***P < 0.001.

For the long-term experiment, the bone marrow cells were harvested from sibling donor mice using the Negative selection EasySep mouse hematopoietic progenitor cell isolation kit (StemCell), following the manufacturer’s instructions. 800 000 enriched HSCs were resuspended in PBS, and IV was injected in the tail vein into irradiated recipient mice while non-irradiated naive mice received PBS. Our previous study demonstrated successful allogeneic transplantation in the bone marrow and spleen six weeks after irradiation using a similar approach^31^. Mice were compared to age-matched sibling controls housed in the same cages.

### Dual-energy X-ray absorptiometry (DXA)

Dual-energy X-ray absorptiometry analysis was conducted several times in the same mice before and after irradiation to evaluate total body areal bone mineral density (aBMD), and lumbar spine (L3–L6) aBMD, using Faxitron UltraFocus DXA (Faxitron Bioptics, LLC, Tucson, AZ, USA). At 40 kV and 0.28 mA for 2.53 seconds, with a spatial resolution of 24 μm and a 2x geometric magnification.

### High-resolution microcomputed tomography (μCT)

High-resolution microcomputed tomography was performed in the femur and lumbar vertebrae (L5) using a Skyscan 1172 scanner (Bruker MicroCT, Aartselaar, Belgium). The lumbar vertebra and femur were imaged with an X-ray tube voltage of 50 kV and a current of 201 μA, with a 0.5 mm aluminum filter, and the scanning angular rotation was 180°, and the angular increment was 0.70°. NRecon (version 1.6.9.8, Bruker MicroCT) was used to reconstruction after scans. In the vertebra, the trabecular bone was analyzed between 235-229 μm from the lower end of the pedicles. In the femur, trabecular bone proximal to the distal growth plate was selected for analyses within a conforming volume of interest (cortical bone excluded), commencing 426 μm from the growth plate and extending a further longitudinal distance of 135 μm in the proximal direction. The cortical measurements in the femur were performed in the diaphyseal region of the femur, starting at 5.23 mm from the growth plate and extending a further longitudinal distance of 135 m in the proximal direction. The data was analyzed using the CTAn software (version 1.13.2.1, Bruker MicroCT). For each time point, age-matched naive littermates were used as the reference point and set to 100%, with the values for the irradiated mice expressed relative to this baseline.

### Peripheral quantitative computed tomography (pQCT)

Trabecular and cortical bone in the femur and tibia were analyzed using the Stratec pQCT XCT Research M (software version 5.4; Norland, Fort Atkinson, WI, USA) at the resolution of 70 μm. The trabecular bone region was defined in the metaphysis area distal to the growth plate in the tibia and proximal to the growth plate in the femur. The area was designated as 2.6% of the length of the bone from the growth plate and only the inner area to 45% of the total cross-sectional area was defined as trabecular bone. Cortical thickness was determined by analyzing scans of the mid-diaphyseal region.

### Murine serum analyses

Blood was collected in blood collection tubes containing a silicon serum separator (Microvette 500 Z-Gel, Sarstedt, Nümbrecht, Germany). Serum levels of total cholesterol in mice were measured using commercial fluorometric assay kits (Sigma-Aldrich CS0005). Enzyme-linked immunosorbent assays (ELISAs) were conducted to quantify C-terminal type I collagen fragments (CTX-I), indicative of bone resorption, (Immunodiagnostic Systems, East Boldon, UK), procollagen type I N pro-peptide (PINP), a serum marker for bone formation (Immunodiagnostic Systems) in serum, and immune cytokines IL-6 (Invitrogen) and TGF-β1 (Invitrogen).

The Proteome Profiler Mouse XL Cytokine Array Kit (R&D Systems) was used to determine the change in serum protein levels following the manufacturer’s instructions. One pooled control serum from naive mice (n=5) was compared to both pooled serum from mice 2 or 12 weeks after irradiation (n=5). The pooled control serum from naive mice served as the baseline and the results are reported as the log2 change in relative units compared to serum from naive mice. We quantified and normalized the signal for the background for each spot using Quick Spots (OptimEyes version 25.5.2.3) and ImageJ (version 1.8.0) software. From the 111 analytes, we selected cytokines, chemokines, and growth factors known from the literature to play a role in skeletal turnover and inflammation. Serum steroid concentrations were measured by high-sensitivity liquid chromatography-tandem mass spectrometry (LC-MS/MS) as described previously^32^.

### Histological examination

Formaldehyde-fixed bone tissue was decalcified in 10% EDTA for 3 weeks. The tibia was then embedded in paraffin and cut into 4 μm thick sections. Hematoxylin and eosin (H&E) staining was performed. Adipocytes were defined as empty white round holes and osteoblast as cubical cells on the bone surface in the H&E staining. TRAP staining was performed for osteoclast detection. In brief, the slides were pre-warmed at 60°C for 30 min and allowed to cool at room temperature. The slides were then washed in Xylene, followed by ethanol and finally with water. Sections were incubated in 0.2 M acetate buffer (pH 5), followed by 1.5 h in TRAP buffer, a solution of Naphthol, red-violet, 0.1 M acetate buffer, 0.3 M sodium tartrate, and Triton X-100, and finally counterstained with fast green and then mounted using Fluoromount Antifade (Sigma-Aldrich). Osteoclasts was defined as TRAP-positive cells on the surface of bone tissue. TUNEL assay (In Situ Cell Death Detection Kit Fluorescein, Roche Diagnostics) was performed according to the manufacturer’s instructions. Briefly, slides were prepared by incubation at 60°C for 60 minutes followed by rehydration through sequential washes in xylene, ethanol (99%, 95%, 70%), and water. The sections were treated with Proteinase K (1:2000 in PBS) at 37°C for 40 minutes in a shaking water bath, followed by washing in PBS (3 × 10 minutes). Labeling was performed using a prepared enzyme and label solution mix, with tissues incubated at 37°C for 1.5 hours under parafilm and light protection. Final washing was done in PBS (3 × 10 minutes).

Images were taken in the epiphyseal and metaphyseal area of the tibia using a Zeiss LSM780 confocal laser scanning microscope with a Zeiss C-Apochromat × 40/1.20 objective (Carl Zeiss). Post-acquisition image processing was performed using Zeiss Zen 3.1 (blue edition) software in the total epiphyseal bone and in a fixed region of interest of 225 μm^2^ 400 μm from the growth plate in the middle region of the bone.

### Gene expression

RNA from cortical bone (tibia), and bone marrow, was extracted using Trizol Reagent (Sigma, St. Louis, MO, USA) followed by RNeasy Mini QIAcube Kit (Qiagen, Hilden, Germany). RNA was reverse transcribed into cDNA using the Applied Biosystems High-Capacity cDNA Reverse Transcription Kit (Applied Biosystems, Waltham, MA, USA). qPCR was run using the StepOnePlus Real-Time PCR systems (Applied Biosystems). Predesigned probes for BAX (Mm00432051_m1) and TGF-β1 (Mm01178820_m1) from Applied Biosystems were used. The mRNA abundance of each gene was calculated using the ΔΔCt method and adjusted for the expression of 18 S ribosomal RNA (4310893E) (Applied Biosystems).

### Cell preparation and flow cytometry

To prepare cells for flow cytometry analysis, bone marrow cells were flushed out and harvested from the femoral bone using a 25 G syringe with PBS. Splenocytes were isolated, and a single-cell suspension in PBS was obtained by passing them through a 70-µm cell strainer. The pelleted cells were resuspended in 0.1 M 1x Tris buffer (pH 7.4) solution to lyse erythrocytes and washed in PBS. The total number of leukocytes was determined using a cell counter (Sysmex, Europe GmBH, Norderstedt, Germany). The fluorochrome-conjugated anti-mouse antibodies were used for flow cytometry analysis: allophycocyanin (APC)-conjugated F4/80, APC-Cy7 conjugated anti-CD8, phycoerythrin (PE)-conjugated anti-CD19 (eBioscience, Thermo Fisher Scientific, Gothenburg, Sweden); APC-cyaninine5(Cy5)-conjugated anti-B220, Brilliant Violet 421-conjugated anti-CD69, V450-conjugated anti-CD11b (Becton Dickinson and Company, Franklin, NJ, USA); and PE-Cy7-conjugated anti-B220, APC-conjugated anti-CD4 and PE-Cy7-conjugated anti-CD8, PerCP-conjugated anti-Gr-1 (eBioscience, Thermo Fisher Scientific, Gothenburg, Sweden), FITC-conjugated Annexin V (Becton Dickinson and Company, Franklin, NJ, USA) and 7AAD (Becton Dickinson and Company, Franklin, NJ, USA). Annexin V and 7AAD staining was performed in 1x Annexin V Binding Buffer containing 0.1M Hepes (pH 7.4), 1.4M NaCl, and 25 mM CaCl_2_ (BD Biosciences). Flow cytometry analyses were performed using FACSVerse (Becton Dickinson) and FlowJo (version 10.6.2) (detailed gating strategies in Fig S5).

### Osteoclast differentiation from primary murine cells or RAW 264.7 cells

Murine osteoclast differentiation was studied in vitro using RANKL-induced differentiation. Bone marrow-derived macrophages (BMMs) were obtained from bone marrow of C57BL/6 mice and cultured in suspension culture dishes (Corning Inc.) with a complete α-MEM medium (Gibco; 12561056), supplemented with 10% heat-inactivated fetal bovine serum (FBS, Sigma-Aldrich; F7524), 100 U/mL penicillin, 100 µg/mL streptomycin (Gibco), and 2 mM GlutaMAX (Gibco). In addition, the medium contained 30 ng/mL macrophage colony-stimulating factor (M-CSF) (R&D Systems). After two days, nonadherent cells were washed away with PBS, and adherent BMMs were detached from the culture dish using 0.02% EDTA in PBS (Sigma-Aldrich). BMMs were then spot seeded in 96-well plates at a density of 5000 cells/5 µL in the center of the wells and incubated with 100ul of media containing 30 ng/mL M-CSF alone or in combination with 4 ng/mL RANKL (R&D Systems; 462-TEC-010) to induce osteoclast differentiation.

The medium was changed or cultivation finished after 3 days when a proportion of the cells had differentiated into multinucleated preosteoclasts or osteoclasts. After three days of cultivation, TGF-β1 protein expression (Invitrogen 88-50690-22) and TRAP5b activity (Bone TRAP, Immunodiagnostic Systems) were measured in the culture supernatant.

RAW 264.7 cells (ATCC; TIB-71) were cultured in complete Dulbecco’s Modified Eagle’s Medium (DMEM, Gibco), supplemented with 10% heat-inactivated fetal bovine serum (FBS, Sigma-Aldrich; F7524), 100 U/mL penicillin, and 100 µg/mL streptomycin (Gibco; 15140122). To induce osteoclast differentiation, RAW 264.7 cells were re-suspended in α-MEM (Gibco), spot-seeded in 96-well plates at a density of 8000 cells/5 µL in the center of the wells and treated with 2 ng/mL RANKL (R&D Systems). The formation of osteoclasts was evaluated by TRAP staining (Sigma-Aldrich) according to the manufacturer’s instructions. Osteoclasts were defined as TRAP-positive cells with three nuclei or more.

### Apoptosis induction

Thymocytes were isolated as apoptotic targets from 4-week-old C57BL/6 female mice, as described before^33^. Cells cultured at a density of 10^7^ cells/mL in α-MEM medium supplemented with 2 mM glutamine, 100 U/mL penicillin, and 100 µg/mL streptomycin. Apoptosis was induced by incubating thymocytes in a serum-free medium for 24 hours to simulate growth factor withdrawal. Freshly isolated thymocytes were used as viable controls for the FACS measurements. The levels of apoptotic and necrotic cells were assessed using 7-AAD/Annexin V staining, and flow cytometry analyses were performed with a FACSVerse flow cytometer (Becton Dickinson) and analyzed using FlowJo software (version 10.6.2) (detailed gating strategies in Fig S6).

### Phagocytosis assay

BMM-derived osteoclasts were prepared as previously described. The cells were spot-seeded at a density of 5,000 cells/5 μL, and after cell attachment, the media volume was adjusted to a total of 200 μL in each well of an 8-well culture slide (Falcon Culture Slides, 354108). The cells were then incubated with 30 ng/mL M-CSF, either alone or in combination with 4 ng/mL RANKL. Thymocytes were labeled with 5 μM Vibrant CFDA (Molecular Probes) for 30 minutes at room temperature and subsequently added to the BMM cultures at a 1:5 cell ratio. Phagocytosis was allowed to proceed either for 60 minutes (short-term assay) or during a 3-day differentiation period (long-term assay) at 37 °C in a 5% CO₂ atmosphere. For the short-term (60-minute) assay, cells were stained with CellMask Orange (Invitrogen) at a final concentration of 2 μg/mL for 15 minutes, and nuclei were counterstained with Hoechst 33342 (Molecular Probes-Invitrogen) at a final concentration of 5 μg/mL, following the manufacturer’s instructions. After the co-culture period, non-internalized thymocytes were thoroughly washed away with PBS, and the cells were fixed with 4% paraformaldehyde (PFA) for 10 minutes at room temperature. Finally, slides were mounted using ProLong Antifade Mountant (Molecular Probes) for imaging. Cells containing two nuclei were classified as pre-osteoclasts, while those containing three or more nuclei were classified as osteoclasts.

### Statistics

All statistical analyses were conducted using GraphPad Prism software (GraphPad Software Inc., La Jolla, CA, USA). All except the sex steroids were normally distributed. Statistical difference in the sex steroids was checked with Mann-Whitney U. A two-sided unpaired Student’s t-test was used to determine statistical differences between two independent groups (naive and irradiated). For multigroup comparisons, one-way ANOVA was performed, followed by Dunn’s test with Šidák correction to compare the control group against each stimulation group (naive versus the different time points post-irradiation). Figures showing data over time are presented as mean ± standard deviation (SD) and significance is indicated as *p <0.05, **p <0.01, ***p <0.001.

## Results

### Total body irradiation has a long-term negative impact on bone mass and uterus weight in female C57BL/6 mice

In the present study, we investigated the effects of total body irradiation followed by hematopoietic stem cell transplantation (HSCT) on female C57BL/6 mice compared to aged-matched littermate naive control mice (Fig. 1A). One week after irradiation, the irradiated and HSCT mice had temporary fluctuations in body weight, after which their weight returned to that of the naive mice (Fig. 1B, Table I). Total and lumbar spine aBMD was measured in the same mice over time, and a significant decrease was observed from four weeks post-irradiation and HSCT compared to naive (Fig. 1C).

**Table 1.** Body and orgain weight Sample sizes ranged from n=8 to 12. Data are presented as mean ± SD. Statistical analysis was performed using the Student’s t-test to assess differences between the irradiated and naive mice at each time point. Significance levels are indicated as *P < 0.05, **P < 0.01, ***P < 0.001

|  |  | Post-irradiation time |  |  |  |
| --- | --- | --- | --- | --- | --- |
|  |  | 1d | 2w | 6w | 12w |
| **Body weight (g)** | **Naive** | 20,64 ± 1,56 | 20,11 ± 2,00 | 21,87 ± 1,92 | 22,31 ± 2,07 |
|  | **Irradiated** | 20,13 ± 1,61 | 18,77 ± 2,18 | 20,93 ± 0,98 | 21,37 ± 1,26 |
| **Spleen /body weight (mg/g)** | **Naive** | 3,05 ± 0,39 | 3,45 ± 0,41 | 3,23 ± 0,34 | 3,58 ± 0,40 |
|  | **Irradiated** | 1,67 ± 0,21*** | 4,01 ± 0,42* | 3,97 ± 0,51** | 3,50 ± 0,31 |
| **Thymus/body weight (mg/g)** | **Naive** | 1,77 ± 0,41 | 2,44 ± 0,30 | 2,02 ± 0,41 | 1,76 ± 0,37 |
|  | **Irradiated** | 0,76 ± 0,18*** | 1,94 ± 0,41** | 2,43 ± 0,47* | 1,61 ± 0,37 |
| **Uterus/body weight (mg/g)** | **Naive** | 2,90 ± 1,30 | 4,27 ± 1,87 | 2,84 ± 1,14 | 2,21 ± 0,58 |
|  | **Irradiated** | 2,48 ± 0,64 | 2,59 ± 0,66* | 0,83 ± 0,28*** | 0,77 ± 0,21*** |

Within a day after irradiation exposure, a significant reduction in spleen and thymus weight was observed (Table I). Two weeks post-irradiation and HSCT, there was a notable increase in spleen weight. In contrast, thymus weight decreased one day after irradiation but returned to levels like the naive mice six weeks post-irradiation and HSCT. Meanwhile, the uterine weight was unchanged the day after irradiation, significant atrophy was noted from the two weeks post-irradiation and HSCT, continuing through the 12-week investigation. No differences were observed in gonadal fat and liver weight (Table I).

Micro-computed tomography (µCT) conducted one-day post-irradiation indicated no direct adverse effects on femoral trabecular bone volume per tissue volume (Tb. BV/TV) (Fig. 1D, 1F). By the second week post-irradiation and HSCT, changes in bone density were observed, including reduced femur Tb. BV/TV, trabecular number, increased trabecular separation, and thickness. These alterations persisted six weeks post-irradiation, except for trabecular thickness. By the 12-weeks mark, the Tb. BV/TV returned to normal compared to controls, but trabecular number and separation changes remained (Fig. 1D, 1F). Complementary pQCT analysis confirmed the µCT findings with a decrease in both tibial and femoral trabecular BMD began two weeks post-irradiation HSCT and persisted throughout the 12-week period (Fig. S1A-B). Furthermore, µCT assessment observed a reduction of the trabecular bone in the axial vertebrae two weeks post-irradiation and HSCT (Fig. S2).

In addition, the cortical thickness in the femur decreased noticeably by two weeks post-irradiation and remained unchanged until 12 weeks, as determined by μCT (Fig. 1G). The same pattern was visualized in the tibia and femur by pQCT analysis, but there was no significant alteration two weeks post-irradiation (Fig. S1A-B).

Mice showed a significant alteration of bone remodeling markers in serum. Bone resorbing marker CTX-I increased 2- and 6 weeks post-irradiation and HSCT (Fig. 1H). Bone formation marker PINP decreased 2 weeks after irradiation, but surprisingly, at 6 weeks, an induction was observed (Fig. 1H).

### Total body irradiation alters bone cell populations and promotes adipogenesis

Following the observed changes in bone density and serum markers of bone remodeling, we examined osteoclasts and osteoblasts in the tibial epiphyseal bone sections (Fig. 2A-B). One day after irradiation and HSCT, there were no immediate changes in the osteoblasts (Fig. 2A). Two weeks after irradiation and HSCT, we observed a reduction in osteoblast surface per bone surface and osteoblast number per bone perimeter, which persisted throughout the observation period. Osteoclast number per bone perimeter declined immediately after irradiation and remained low through the following six weeks, while osteoclast surface per bone surface only differed at 6 weeks after irradiation (Fig. 2B). Adipocytic expansion within the bone marrow in both metaphyseal as well as epiphyseal bone 2- and persists until 12-weeks post-irradiation and HSCT (Fig. 2C-D).

**Fig. 2:**
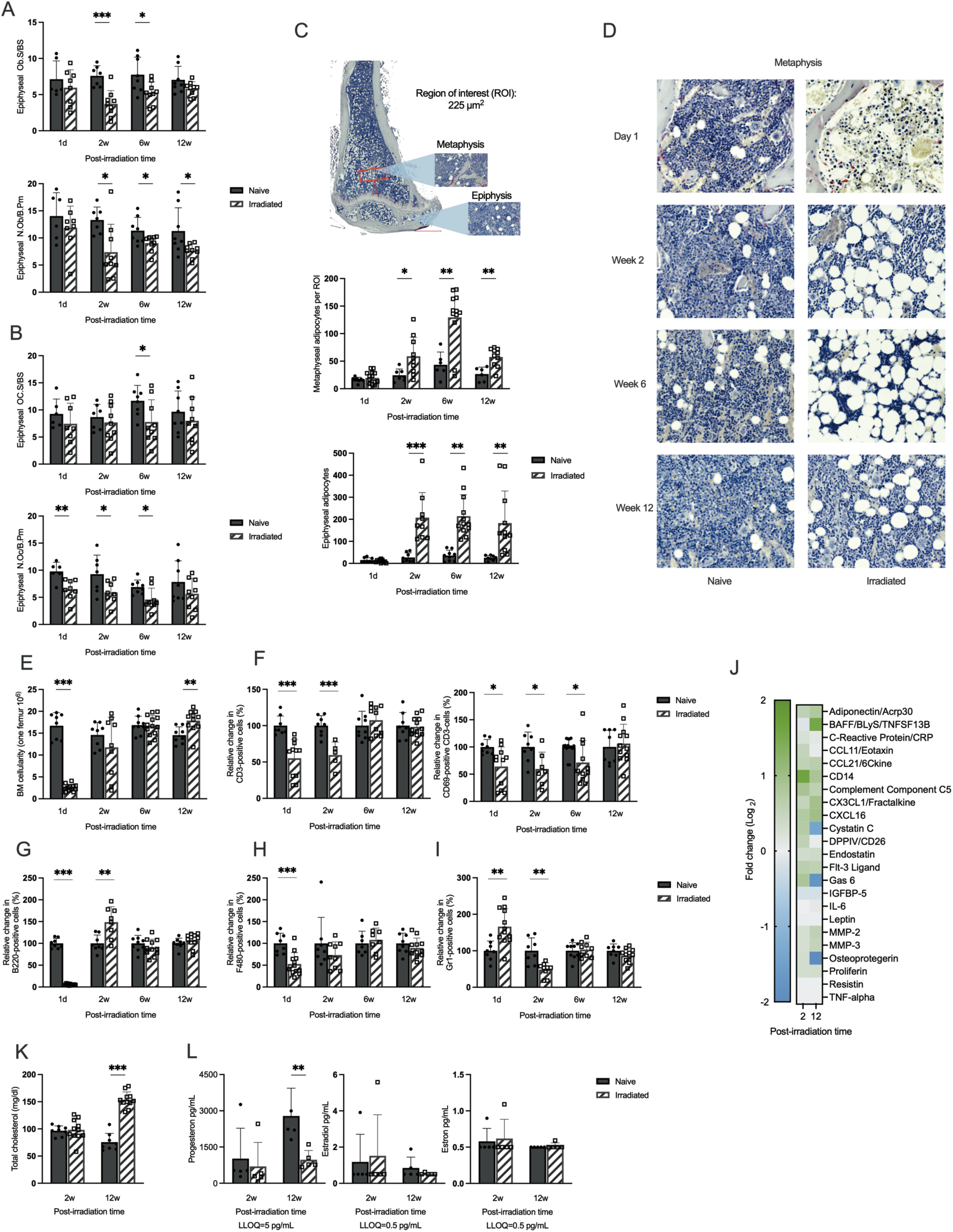
Bone turnover and immune parameters in response to radiation. **A**. Osteoblast surface per bone surface (Ob S/BS) and number of osteoblasts per bone perimeter (N Ob/B Pm) in the distal epiphyseal part of the tibia. **B**. Osteoclast surface per bone surface (Oc S/BS) and number of osteoclasts per bone perimeter (N Oc/B Pm) in the distal epiphysis of the tibia. **C**. Quantification of bone marrow adipocyte number in a region of interested in metaphyseal and epiphyseal of the tibia. **D**. Hematoxylin and eosin staining of the tibia to observe cellular changes. **E**. Number of bone marrow cells per femur. **F**. Relative change in frequency of CD3^+^ T cells and CD69^+^ activated T cells gated on the CD19^-^ cell population. **G**. Relative change in frequency of B220^+^ B cells gated on the CD3^-^ cell population. **H**. Relative change in frequency of F480^+^ monocytic subset gated from the CD3^-^CD11b^+^ population. **I**. Relative change in frequency of Gr1^+^ neutrophil cells gated from the CD3^-^CD11b^+^ population. **J**. Protein composition in serum post-irradiation and HSCT was detected by Mouse Cytokine microarray membrane analysis. Log_2_ fold change was calculated using the corrected pixel density. **K**. Comparative analysis of total serum cholesterol levels. **L**. Measurement of female sex steroids progesterone, estradiol, and estrone levels in the serum (n=5). Statistical analysis was performed using Student’s t-test to assess differences at each time point. Except in sex steroids where Mann-Whitney was performed do to not normally distributed values. The number of samples ranged from n=5-12. Data are presented as mean ± SD. Significance levels are indicated as *P < 0.05, **P < 0.01, ***P < 0.001.

A distinct reduction of bone marrow cells was observed in the histological section one day after irradiation (Fig. 2D, top panels). This reduction was confirmed by counting the cellularity in the femur (Fig 2E). The cellularity was restored at 2- and exceeded the naive mice cellularity 12-weeks post-irradiation and HSCT (Fig. 2E). Not only absolute numbers but also the frequency of several bone marrow cell types, such as CD3^+^ T cells, activated CD69^+^ T cells, B220^+^ B cells, and the F480^+^ macrophage subset, decreased one day after irradiation (Fig. 2F-H). Conversely, the frequency but not the absolute numbers of Gr1^+^ neutrophils rapidly increased one day after irradiation but decreased two weeks after irradiation and HSCT (Fig. 2I). The frequency of CD3^+^ T cells remained lower than the naive mice two weeks after irradiation and HSCT. By 12 weeks post-irradiation and HSCT, the frequencies of all investigated cell types had returned to normal levels. This notable restoration of cell frequencies over time was also observed in the spleen, where a similar rapid decrease in lymphocytes and elevated levels of B cells at 2 weeks after irradiation (Fig. S3A-D). The elevated Gr1^+^ neutrophil population was also observed in the spleen, which returned to similar levels observed in naive mice six weeks post-irradiation and HSCT (Fig. S3A-D).

To analyze the differential protein expression changes affecting systemic bone turnover and immune regulation, we performed a semi-quantitative microarray on the serum, comparing 2- and 12-week post-irradiation and HSCT. The serum levels of B-cell activating factor (BAFF) were indicated elevated, while the levels of cystatin C, growth arrest-specific protein 6 (GAS6), and osteoprotegerin (OPG) were decreased at the 12-week time point compared to the naive mice (Fig 2J, Table S1). Interestingly, key inflammation markers, including IL-6, remained unaffected, confirmed with ELISA (Fig 2J, Fig S3E). Total serum cholesterol levels increased 12 weeks after irradiation (Fig. 2K). In addition, serum progesterone levels decreased 12 weeks post-irradiation and HSCT, but estradiol levels were under the detection limit in both naive and irradiated mice (Fig. 2L).

### Persistent TUNEL-positive cells detected in the bone post-irradiation

As expected, the bone marrow cellularity dramatically decreased one day after irradiation (Fig. 2D, E). Flow cytometry analysis revealed increased apoptotic cells at this time point but no change in the necrotic cells (Fig. 3A). TUNEL staining confirmed this apoptotic cell presence in epiphysis and metaphysis bone marrow as well as within the bone tissue of the tibia one day after irradiation and surprisingly persisted during 12-weeks investigated time (Fig. 3B). The number of TUNEL positive cells were induced 2- and 6-weeks post-irradiation and HSCT but present also 12-weeks after. The apoptotic cells in bone marrow were also accompanied by an upregulation of pro-apoptotic BAX gene expression in the bone marrow and cortical bone one-day post-irradiation and remained elevated for 6 weeks in the bone marrow and 2 weeks in cortical bone (Fig. 3C). In addition, the gene expression of TGF-β1 was elevated after irradiation in bone marrow, but not in cortical bone (Fig. 3D). Furthermore, TGF-β1 in serum was borderline elevated one-day after irradiation (Fig S3F).

**Fig. 3:**
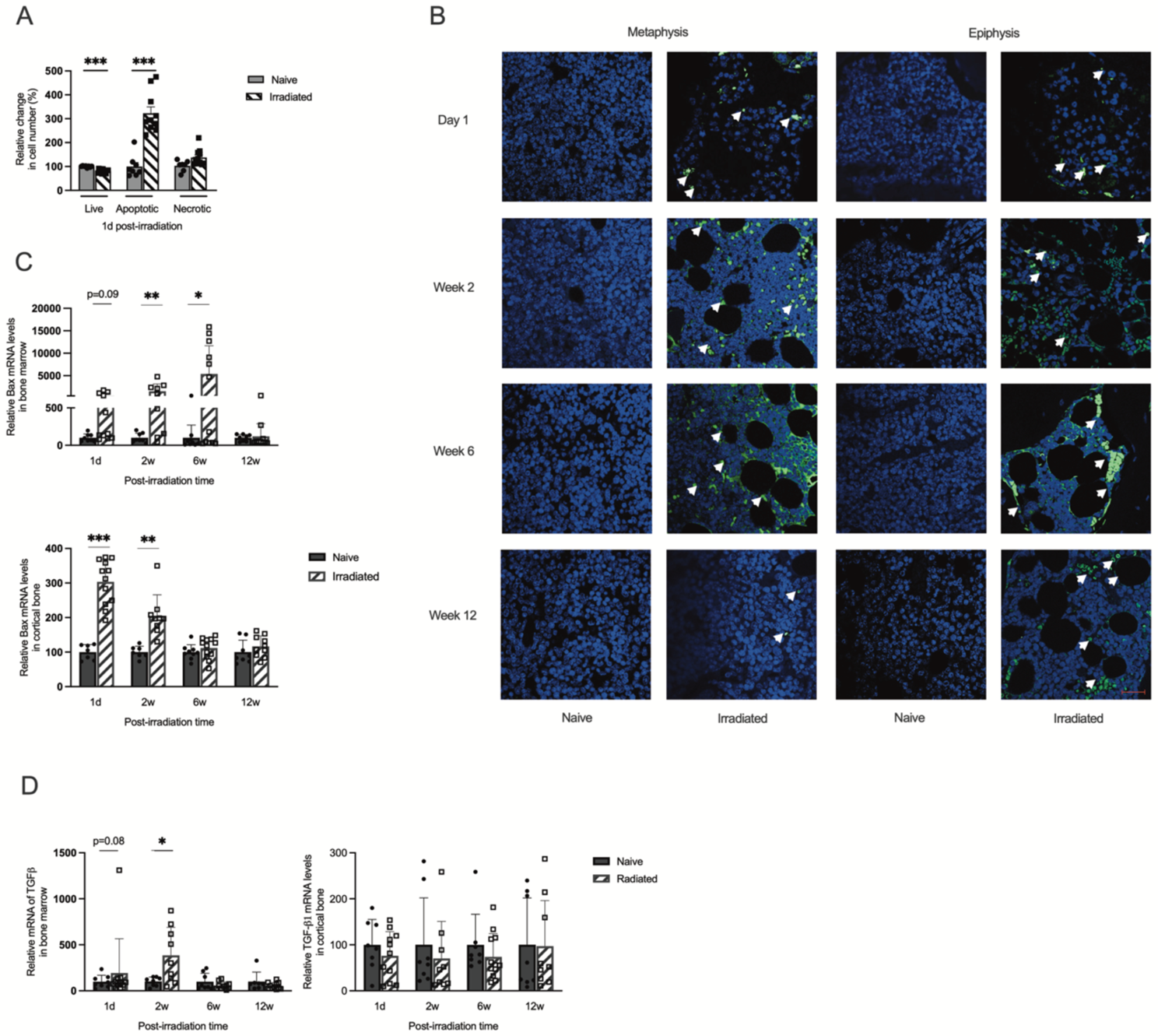
Characterization of radiation-induced cell death. **A.** Flow cytometric detection of relative change in apoptosis and necrosis in bone marrow cells one day post-radiation, utilizing Annexin V-FITC/7-AAD staining. **B.** Representative images of TUNEL staining in the femur bone marrow and epiphysis. All images are captured at 400x magnification with a scale bar of 40 µm. **C.** Analysis of relative gene expression of Bax gene in bone marrow and cortical bone. **D.** Analysis of relative gene expression level of TGF-β1 gene in bone marrow and cortical bone. The number of samples ranged from n=8-12. Data are presented as mean ± SD. Statistical significance was determined using Student’s t-test at each time point, with p-values indicated as *P < 0.05, **P < 0.01, ***P < 0.001.

### Total body irradiation showed no immediate effect on bone microarchitecture but altered the bone marrow cells

To further investigate the immediate changes after irradiation that may develop into long-term effects, we conducted an experiment using a single irradiation of 9 Gy to elucidate the underlying mechanisms (Fig. 4A). In response to irradiation, body weight remained unchanged during the first 48 hours (Table II). Femur trabecular bone mineral density and cortical thickness determined by pQCT were neither altered within this limited timeframe (Fig. 4B).

**Fig. 4:**
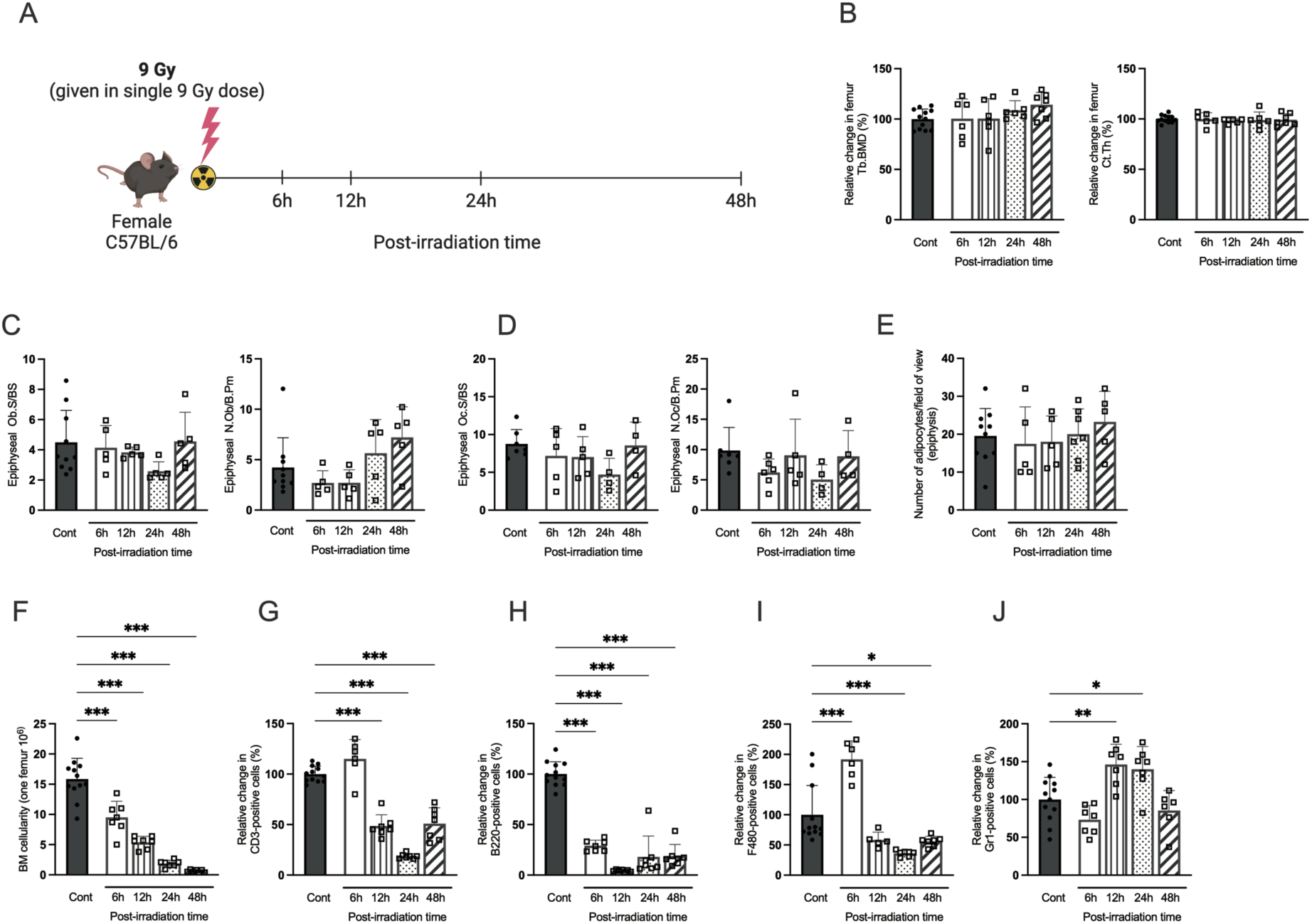
Acute effect of the radiation on bone parameters. **A.** Experimental design of the acute radiation scheme. **B**. Peripheral Quantitative Computed Tomography (pQCT) measurement of trabecular Bone Mineral Density (Tb.BMD) and cortical thickness (Ct.Th) in irradiated and naive mice. **C**. Number of osteoblasts per bone perimeter (N Ob/B Pm) and osteoblast surface per bone surface (Ob S/BS) in the distal epiphyseal part of the tibia. **D**. Number of osteoclasts per bone perimeter (N Oc/B Pm) and osteoclast surface per bone surface (Oc S/BS) in the distal epiphysis of the tibia. **E**. Quantification of bone marrow adipocyte number epiphyseal of the tibia. **F**. Number of bone marrow (BM) cells per femur. **G**. Relative change in CD3^+^ T cells gated on the CD19^-^ cell population. **H**. Relative change in B220^+^ B cells gated on the CD3-cell population. **I**. Relative change in F480^+^ monocytic subset gated from the CD3^-^CD11^+^ population. **J**. Relative change in Gr1^+^ neutrophil cells gated from the CD3^-^CD11^+^ population. For statistical analyses, one-way ANOVA and Dunnett’s multiple comparisons test were used to compare the means of a control group with each time point. The number of samples ranged from n=8-12. Data are presented as mean ± SD., and significance is indicated as *P < 0.05, **P < 0.01, ***P < 0.001.

**Table 2.** Body and orgain weight Sample sizes ranged from n=8 to 12. Data are presented as mean ± SD. Statistical analysis was performed using the Student’s t-test to assess differences between the irradiated and naive mice at each time point. Significance levels are indicated as *P < 0.05, **P < 0.01, ***P < 0.001

|  |  | Post-irradiation time |  |  |  |
| --- | --- | --- | --- | --- | --- |
|  | Naive | 6h | 12h | 24h | 48h |
| **Body weight (g)** | 20,11 ± 0,98 | 18,99 ± 0,80 | 19,89 ± 0,84 | 18,99 ± 1,67 | 19,20 ± 1,17 |
| **Spleen/body weight (mg/g)** | 3,13 ± 0,37 | 2,49 ± 0,38*** | 2,04 ± 0,35*** | 1,59 ± 0,15*** | 1,41 ± 0,25*** |
| **Thymus/body weight (mg/g)** | 2,35 ± 0,55 | 2,00 ± 0,22 | 1,65 ± 0,15*** | 0,87 ± 0,17*** | 0,42 ± 0,11*** |
| **Uterus/body weight (mg/g)** | 2,93 ± 1,30 | 3,06 ± 1,28 | 3,24 ± 0,83 | 2,85 ± 2,05 | 1,62 ± 0,10 |
| **Liver/body weight (mg/g)** | 42,24 ± 3,46 | 39,77 ± 3,95 | 37,15 ± 2,05** | 43,32 ± 3,48 | 42,83 ± 2,27 |

Reduction in spleen weight was noted in the irradiated group as early as 6 hours post-irradiation, with the decrease continuing up to 48 hours (Table II). Thymus weight was reduced 12 hours post-irradiation, with continuing decreases up to 48 hours. Significant alterations in liver weight were observed only 12 hours after irradiation. Uterine weight remained unaffected. No changes were observed in the numbers of osteoblasts, osteoclasts, and adipocytes, nor the size of osteoclasts and osteoblasts in the tibial epiphyseal region (Fig. 4C–E). Bone marrow cellularity gradually decreased in the irradiated group compared to the naive (Fig. 4F). The frequency of the CD3^+^ T cell population decreased 12 hours post-irradiation (Fig. 4G), while the B220^+^ B cell population showed greater radiosensitivity as its frequency dramatically decreased immediately after irradiation (Fig. 4H). For the F480^+^ monocytic population, a rapid increase in cell frequency was observed 6 hours after irradiation, then decreased at 24- and 48 hours post-irradiation (Fig. 4J). Finally, the Gr1^+^ neutrophil population increased 12- and 24 hours post-irradiation (Fig. 4K). This same pattern of alteration in cell frequencies was also observed in the spleen, with a similar rapid decrease in lymphocytes and an increment of neutrophils and monocytes (Fig. S4).

Flow cytometry analysis showed that most cells in the bone marrow of naive mice were viable (Fig. 5A). Following irradiation, the frequency of viable cells decreased following the reduced number of bone marrow cells, while apoptotic cells increased up to 48 hours post-irradiation. TUNEL staining showed dying cells in the bone tissue of both the epiphyseal and metaphyseal regions as early as 6 hours post-irradiation, persisting across all subsequent time points (Fig. 5B).

**Fig. 5:**
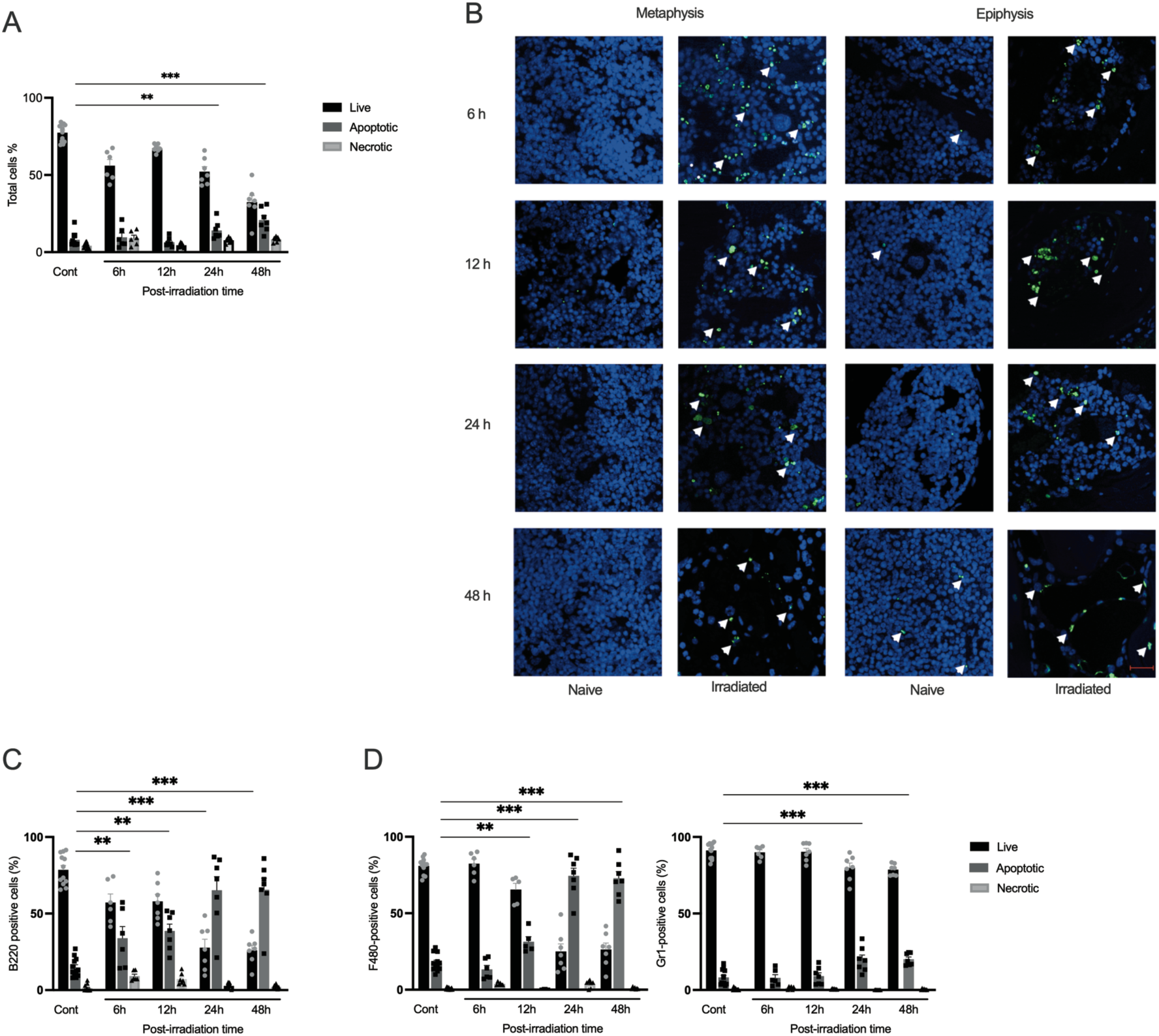
Acute bone and bone marrow injury within 48 hours post-radiation. **A**. Percentage of live, apoptotic, and necrotic cells measured by flow cytometry using Annexin V-Fitc/7-AAD staining assay in the total bone marrow cell population. **B**. Representative images of TUNEL staining of the tibia in metaphysis and epiphysis. **C**. Percentage of live, apoptotic, and necrotic cells in B220^+^ B cells gated on the CD3^-^cell population. **D**. Percentage of live, apoptotic, and necrotic cells in F480+ monocytic subset and Gr1^+^ neutrophil cells gated from the CD3^-^CD11^+^ population. Apoptotic cells were identified as Annexin V+ (Annexin V+), necrotic cells as both Annexin V and 7-AAD+ (Annexin V+ 7AAD+), live cells as negative for both Annexin V and 7-AAD (Annexin V− 7AAD−), and cell fragments or debris were characterized by high 7-AAD density. Statistical analysis involved one-way ANOVA followed by the Dunn–Šidák post-test. Data are presented as mean ± SD, and significance is indicated as *P < 0.05, **P < 0.01, ***P < 0.001 compared to the control live population.

Within 6 hours post-irradiation, B cells showed a sharp increase in apoptosis, peaking at 24 hours (Fig. 5C). monocytes showed a significant rise in apoptosis at 12 hours (Fig. 5D). Neutrophils displayed only a limited but significant increase in apoptotic response 24- and 48-hours post-irradiation (Fig. 5E).

### Pre-osteoclasts can engulf apoptotic cells during in-vitro differentiation

Apoptotic cells in the bone marrow compartment during the 12-week follow-up after irradiation may influence bone resorption. Mimicking this effect, apoptotic lymphocytes from thymuses were added to osteoclast cultivation *in vitro*. Primary murine bone marrow macrophage (BMM) and RAW264.7 cells were differentiated with RANKL to induce osteoclast differentiation or maintained as macrophages as a control. By day 3 of the osteoclast culture, TRAP-positive osteoclasts with more than three nuclei, preosteoclasts with two nuclei were formed, and mononuclear undifferentiated progenitors were observed (Fig. 6A-B Fig. S6A-B).

**Fig. 6.**
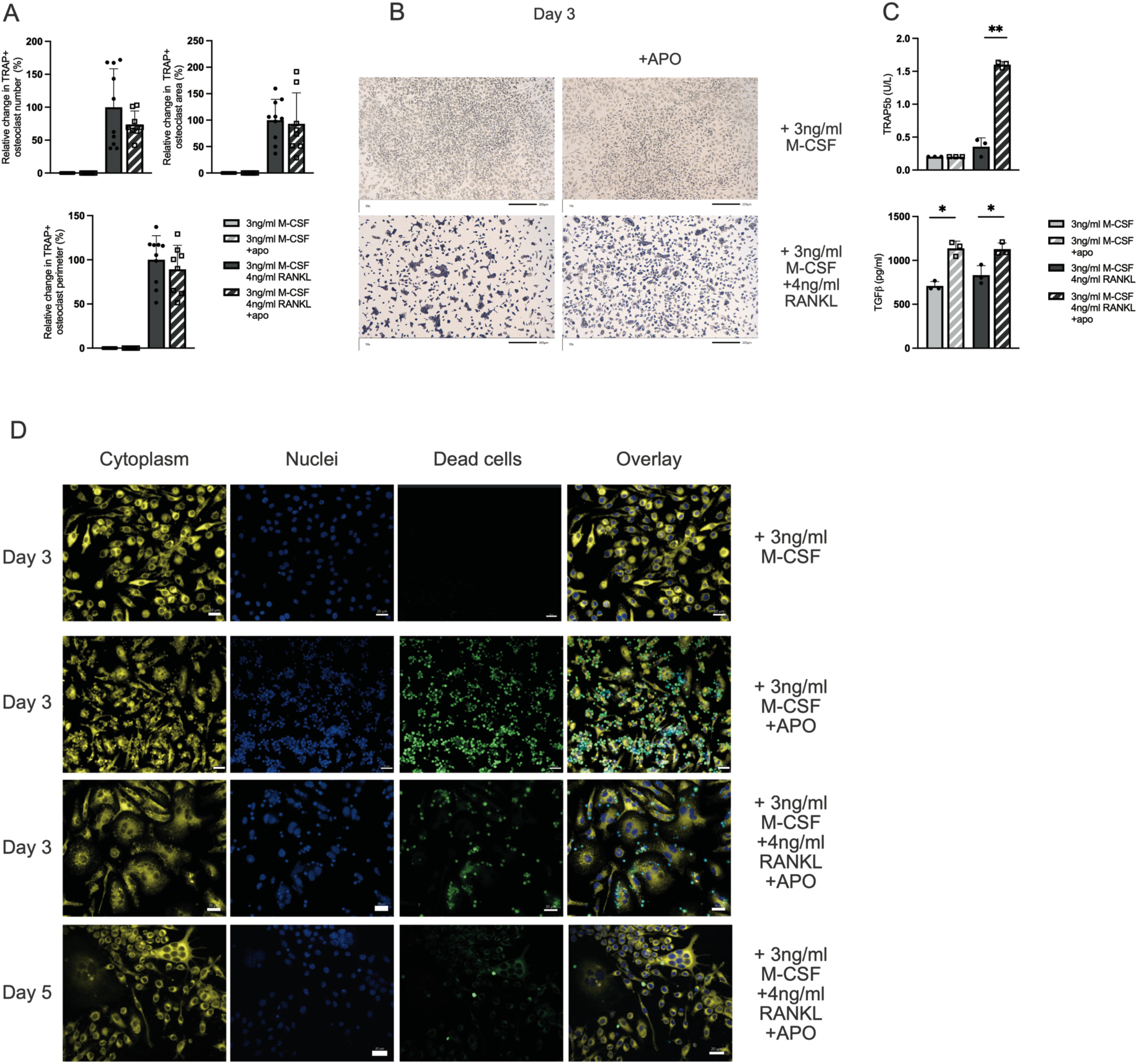
Apoptotic cells did not influence osteoclast morphology in bone marrow macrophages (BMM) during *in vitro* differentiation but initiated osteoclast phagocytosis. **A**. Relative difference changes in the number, area, and perimeter of multinucleated TRAP+ cells after 3 days of differentiation, with or without apoptotic (APO) cells. 4 independent experiments with total 11 mice. **B**. Representative images of osteoclast differentiation after 3 days of stimulation with or without the presence of apoptotic cells. **C**. TRAP5b and TGF-β protein expression in the supernatant of osteoclast differentiation (n=4). **D**. Microscopic observation of the osteoclast phagocytosis assay Statistical analysis was performed using the unpaired t-test. Data are presented as mean ± SD, and significance is indicated as *P < 0.05, **P < 0.01.

When apoptotic cells were added to the BMM culture, they were evenly distributed across the well (Fig. S6C). After three days, BMM macrophages cocultured with apoptotic cells showed significant engulfment. A higher number of apoptotic cells remained along the border of the well compared to the center, where the cells were spot-seeded. When RANKL was added to drive osteoclast differentiation, the engulfment of the apoptotic cells was observed, but to a limited degree compared to the control macrophages. Apoptotic cells did not influence osteoclast number, area, or perimeter (Fig. 6A-B, Fig S6A-B). However, adding apoptotic cells increased TRAP5b levels in the supernatant 3-days in the BMM osteoclast differentiation (Fig. 6C). Interestingly, in the same supernatant, we observed an increased level of TGF-β1 after adding apoptotic cells both in macrophages and in the cells pushed to become osteoclast with RANKL.

After three days of differentiation with M-CSF, apoptotic cells were incubated for 60 minutes with BMM. During this time, phagocytosis was observed, with apoptotic cells found inside the macrophage cytoplasm (Fig. 6D). In cultures treated with RANKL for three days to induce osteoclast differentiation, both pre-osteoclasts and osteoclasts engulfed apoptotic cells. Interestingly, fully matured osteoclasts with more than three nuclei showed a reduced ability to engulf apoptotic cells 3- and 5-days after RANKL treatment (Fig. 6D).

## Discussion

Although extensive research has established that total body irradiation (TBI) disrupts bone homeostasis, certain aspects of immune function, and the bone marrow niche, much of the existing literature has focused on acute or short-term effects. Studies have previously shown that irradiation-induced bone loss could be mediated by bone mesenchymal stem cell dysfunction^2^, increased senescent in both mesenchymal as well as hematopoietic cells^34^, or an increase of hypoxic bone microenvironment^35^. While these processes are likely central to irradiation-induced bone loss, we also believe that the apoptotic cells may critically influence osteoclasts and bone health. Our study explored the effects of irradiation focusing on bone mineral density and bone marrow in female C57BL/6 mice both in an acute phase, up to 48 hours after irradiation, and long-term up to 12 weeks after irradiation and hematopoietic stem cell transplantation (HSCT). By examining these outcomes in both the acute and extended phases post-irradiation, our study offers insights into the persistent consequences of irradiation and HSCT on bone and bone marrow homeostasis.

In the acute phase, a transient increase in neutrophil frequency one-day post-irradiation indicated an inflammatory response, but all other immune cells showed marked reductions. Focusing on compensatory induction of immune function revealed significant increases in spleen weight and B cell frequencies two weeks after irradiation and HSCT, accompanied by increased serum BAFF levels. This suggests that spleen weight recovery depends partly on the repopulation of hematopoietic cells, particularly B cells. In contrast, thymus weight and T cell frequencies remained reduced at two weeks post-irradiation and HSCT but normalized by six weeks, indicating a direct association between thymus atrophy and T cell depletion.

Bone marrow cellularity normalized two weeks post-irradiation and HSCT and increased above the naive levels by twelve weeks, reflecting a compensatory proliferation. However, this increase in absolute immune cell numbers was not mirrored by changes in cytokine expression in bone marrow or protein levels in serum, suggesting a localized rather than systemic immune response to irradiation.

The uterus showed progressive atrophy two weeks post-irradiation, likely linked to the previously observed high radiosensitivity^36,37^. This atrophy in the uterus was further confirmed with decreased serum levels of progesterone, but estrogen and estradiol levels were undetectable in naive and irradiated HSCT mice. Cholesterol, a metabolic marker and the precursor for the sex steroids was elevated twelve weeks post-irradiation, pointing to alterations that require further investigation.

The high calcium content in bone makes it radiosensitive^3,4^. In addition, the bone is a specific environment where hematopoietic and mesenchymal cells regulate each other, and there is evidence that hematopoietic-lineage cells are intertwined with bone tissue^38^. Irradiation significantly disrupted both trabecular and cortical bone parameters. Total areal bone mineral density declined after four weeks, and this decline persisted for up to twelve weeks post-irradiation and HSCT. Cortical thickness reduction and trabecular microarchitecture reductions were evident as early as two weeks, characterized mainly by reduced trabecular number and increased separation. These long-term structural impairments underline the chronic impact of irradiation on skeletal health.

Elevated CTX-I levels (a marker of bone resorption) and an initial decrease in PINP levels (a marker of bone formation) indicated a shift toward reduced bone. The PINP levels increased by six weeks, suggesting a possible adaptive recovery phase in bone formation. Together with the semi-quantitative microarray, which indicated reduced serum osteoprotegerin as an inhibitor of osteoclast activity, it highlights continued dysregulation of the osteoclast activity.

Osteoblast numbers and the surface covering bone area declined two weeks post-irradiation. While osteoclast numbers decreased immediately following irradiation, indicating higher sensitivity to irradiation-induced cell death^39,40^. Indeed, no alteration was observed in the bone area cover of osteoclasts, indicating that the osteoclast gets bigger and still covers the same area. Furthermore, the increase in bone marrow adipocytes indicates a shift in mesenchymal differentiation from osteogenic to adipogenic lineage^41–43^, likely impairing bone regeneration following irradiation-induced damage^44^. This was consistent with an indication of reduced RUNX2 expression, a key regulator of osteoblast differentiation. The mechanisms driving adipocyte induction and whether this impacts the osteoblast formation in this context remain to be fully explained.

A critical observation was the sustained presence of apoptotic cells in the bone marrow and bone tissues up to twelve weeks post-irradiation. TUNEL-positive cells and elevated expression of the pro-apoptotic gene BAX persisted, suggesting prolonged apoptosis and insufficient efferocytosis. This was accompanied by increased TGF-β1 gene expression in the bone marrow during the first two weeks post-irradiation and HSCT. TGF-β1 which stimulated osteoclast, is one of the most abundant cytokines in the bone matrix^45^ is released during bone resorption^46^, and is known to be induced by radiation^47,48^ and efferocytosis^49^. Our experiments did not observe significant changes in cortical bone TGF-β1 gene expression and only limited increases in serum TGF-β1 levels, suggesting a localized regulatory effect confined to the bone marrow.

In the acute irradiation approach, it was visible that the lymphocytes are the most radiosensitive cells affected most rapidly into apoptosis. Macrophages were affected within the first two days, while neutrophils exhibited only a limited response. Apoptotic bone lining cells and osteoblasts from microfractures are known to stimulate osteoclasts by releasing cytokines and TGF-β1^50^. Interestingly, in our *in vitro* experiments, TGF-β1 protein levels were elevated in the supernatant from osteoclast differentiation after a 3-day co-incubation with apoptotic cells from lymphocytes compared to cultures without apoptotic stimuli. In contrast, it is well established that macrophages engulfing apoptotic cells can produce TGF-β1^51–53^. Our findings indicate that apoptotic stimuli during osteoclast differentiation can maintain the TGF-β1 production. This can potentially contribute to TGF-β1 expression in the bone marrow, which could play a role in modulating bone remodeling.

Prolonged exposure to apoptotic cells did not affect the osteoclast number or morphological parameters in murine osteoclasts, a confirmed model for *in vitro* osteoclast differentiation^54–56^. Indeed, our experiments observed elevated TRAP5b protein levels in the supernatant after three days of osteoclast differentiation in the presence of apoptotic cells, compared to cultures without apoptotic stimuli. Although higher TRAP5b activity has been associated with an increased osteoclast number *in vitro*^57^, no difference in osteoclast numbers could be detected in our studies.

Pre-osteoclasts and smaller osteoclasts rapidly engulfed apoptotic cells within 60 minutes, demonstrating efferocytosis activity, though this did not influence the number of mature osteoclasts. Indeed, previous studies have shown that osteoclasts activated as antigen-presenting cells maintain their bone resorbing capacity^17^. The role of osteoclasts in efferocytosis was first described by Harre et al., who demonstrated that human osteoclasts can engulf apoptotic cells and express genes associated with the efferocytosis machinery^19^. The efferocytosis in bone was further described by Batoon et al., who described how affecting the apoptotic capacity in mice influences bone tissue^20,22^. Our findings demonstrate that murine bone marrow-derived pre-osteoclasts stimulated for three days with M-CSF and RANKL exhibit efferocytosis activity. In contrast, this capability was reduced in fully matured osteoclasts. Increased TGF-β1 expression during *in vitro* cultivation, secreted during efferocytosis, modulating osteoclast activity and influencing the bone microenvironment. The sustained population of apoptotic cells in bone marrow after irradiation and bone marrow transplantation likely contributes to local stimulation of bone-resorbing osteoclasts and increased adipocyte numbers, collectively impairing bone regeneration.

While irradiation does not cause immediate bone loss, it leads to a continuous decline in cortical and trabecular bone beginning two weeks post-irradiation and persisting throughout our twelve-week study duration. Promoting efficient efferocytosis in the bone could mitigate these effects and offer a therapeutic strategy to improve skeletal health.

## Ethics statement

The study was approved by the ethics committees of the Gothenburg region, Sweden (case number 1–2017).

## Supporting information

all supp

## Acknowledgments

The authors gratefully acknowledge the valuable technical assistance and contributions of Jianyao Wu, Anna-Karin Norlén, Ifrah Samater, Vilma Enlund, Zhicheng Hu, Aidan Barret, and Catarina Magnusson.

## Authors contributions

Conceptualization, T.S. and C.E: Formal analysis, T.S., P.G., and C.E; Animal experimentation, T.S., P.G., K.H., P.H., M.K.L, and C.E; Histological assessment, T.S., C.C., A.S.C., and C.E; Bone assessment, C.O. and M.K.L; Sex steroids measurement C.O. Cell cultivation, T.S., and C.E; Responsible for acquisition of data: T.S., P.G., A.S.C., and C.E; Funding acquisition, C.E., A.S.C.; Writing—original draft: T.S. and C.E: Writing—review and editing: T.S., P.G., K.H., P.H., C.C., M.K.L, A.S.C., and C.E.

