## Supplementary material for "Disrupted bone microenvironment and immune recovery following total body irradiation in a murine model": all supp

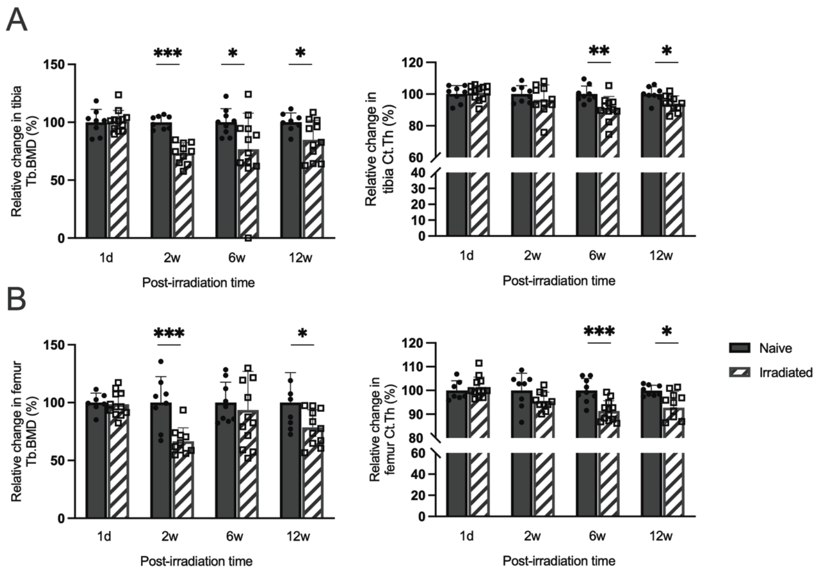


**Fig. S1: pQCT analysis of tibia and femur.**

**A**. Mice body weight at each sacrifice time point. **B**. Trabecular volumetric bone mineral density (Tb. BMD) and cortical bone thickness (Ct. Th.) measured in the tibia by Peripheral Quantitative Computed Tomography (pQCT). **C**. Trabecular volumetric BMD and cortical bone thickness measured in the femur by pQCT. Statistical analysis was performed using the Student's t-test to assess differences between the irradiated and control mice at each time point. Sample sizes ranged from n=8 to 12. Data are presented as mean ± SD. Significance levels are indicated as *P < 0.05, **P < 0.01, ***P < 0.001.

Sup fig 2

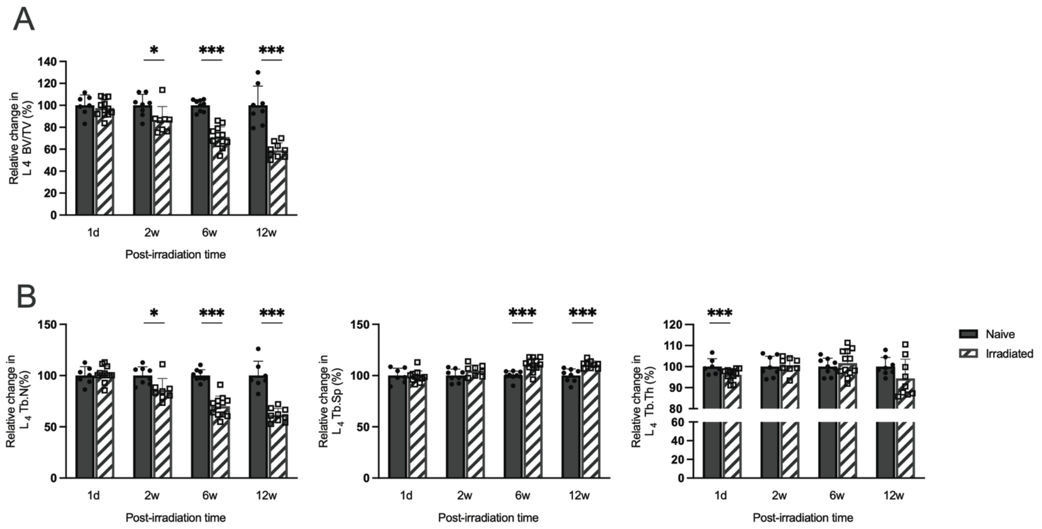


**Fig. S2: High-resolution microcomputed tomography (µCT) analysis of L4 vertebrae.**
**A**. Relative change in micro-computed tomography (µCT) analysis of trabecular bone volume per total volume (Tb. BV/TV) in lumbar vertebra 4 (L4). **B**. Relative change in L4 Trabecular Number (Tb.N), Trabecular Separation (Tb.Sp), and Trabecular Thickness (Tb.Th). Statistical analysis was performed using the Student's t-test to assess differences between the irradiated bone marrow transplanted and control mice at each time point. Sample sizes ranged from n=8 to 12. Data are presented as mean ± SD. Significance levels are indicated as *P < 0.05, **P < 0.01, ***P < 0.001.

Sup fig 3

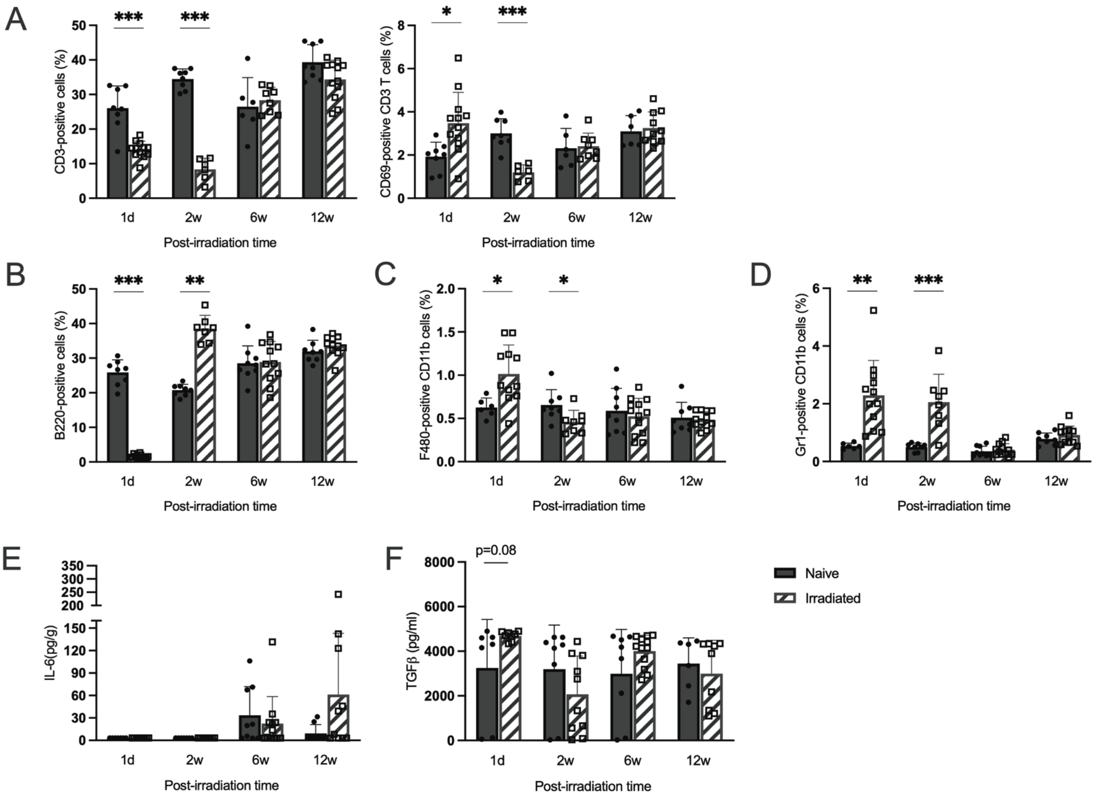


**Fig. S3: Flow cytometric analysis of the spleen cell population in irradiated and bone marrow transplanted mice.**

**A**. Relative change in $CD3^+$ T cells gated on the $CD19^-$ cell population. **B**. Relative change in $B220^+$ B cells gated on the $CD3^-$ cell population. **C**. Relative change in $F480^+$ monocytic subset gated from the $CD3^-CD11b^+$ population. **D**. Relative change in $Gr1^+$ neutrophil cells gated from the $CD3^-CD11b^+$ population. **E**. IL-6 protein levels were quantified in the irradiated and HSCT mice serum and compared to control naive mice. **F**. TGF-β1 protein levels were quantified in the serum of irradiated mice and compared to naive (non-irradiated) mice. Statistical analysis was performed using the Student's t-test to assess differences between the irradiated bone marrow transplanted mice and control mice at each time point. Data are presented as mean ± SD. Significance levels are indicated as *P < 0.05, **P < 0.01, ***P < 0.001.

Sup fig 4

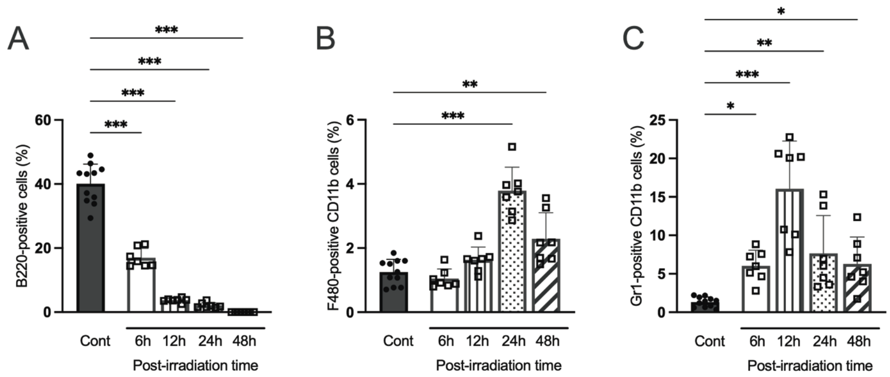


**Fig. S4: Flow cytometric analysis of the spleen cell population in irradiated and bone marrow transplanted mice.**

**A**. Relative change in B220$^+$ B cells gated on the CD3$^-$ cell population. **C**. Relative change in F480$^+$ monocytic subset gated from the CD3$^-$CD11$^+$ population. **D**. Relative change in Gr1+ neutrophil cells gated from the CD3$^-$CD11$^+$ population.

Sup fig 5

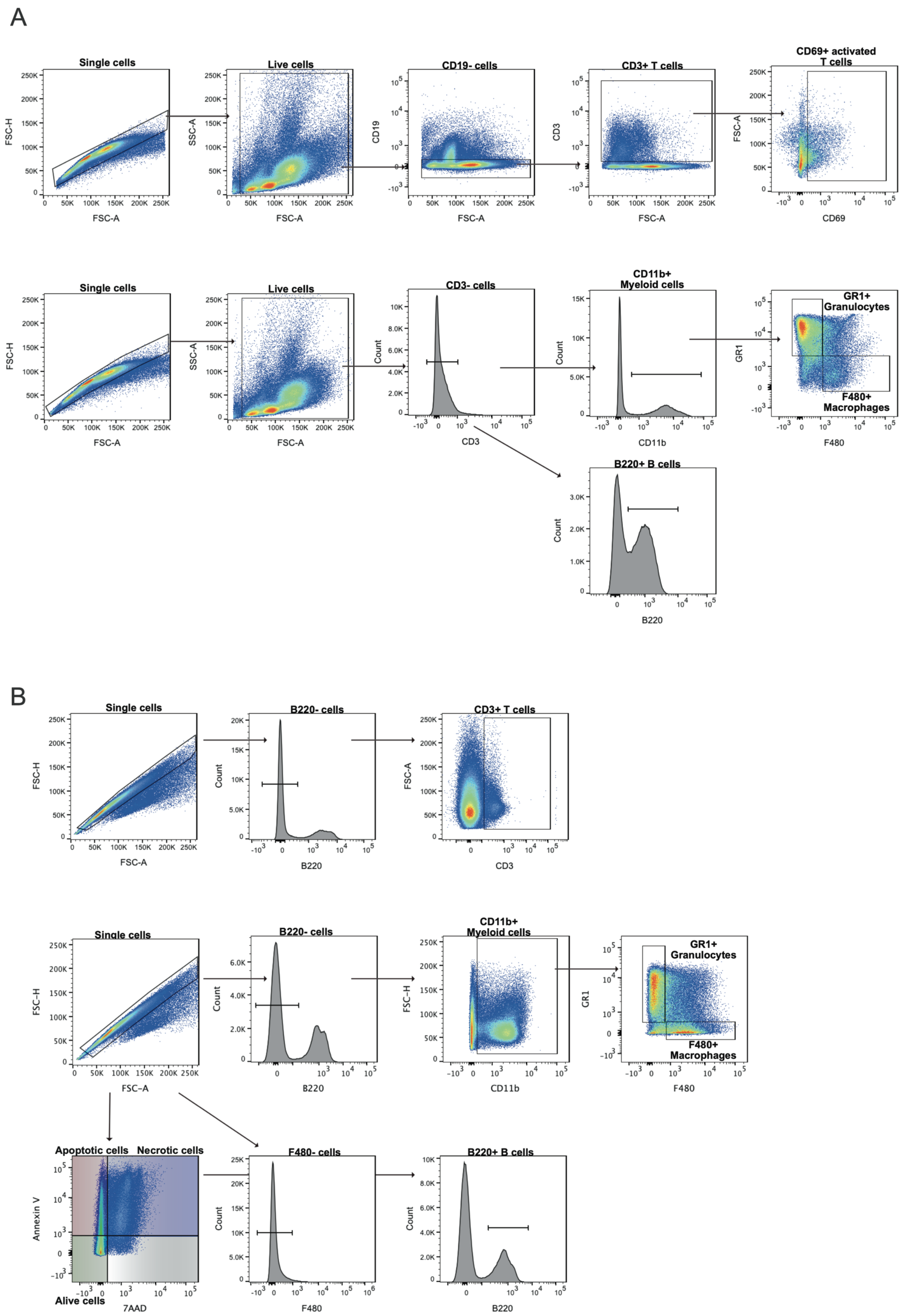


**Fig. S5: Representative gating strategies for the bone marrow and spleen cell populations.**

**A**. Cells from bone marrow were gated for forward scatter height vs. forward scatter area (FSC-H/FSC-A) before identification of total T cells (CD19$^-$CD3$^+$), activated T cells (CD19$^-$CD3$^+$CD69$^+$), granulocytes (CD3$^-$CD11b$^+$Gr1$^+$), macrophages (CD3$^-$CD11b$^+$F4/80$^+$), and pan B cells (CD3$^-$B220$^+$). **B**. Cells from bone marrow were gated for forward scatter height vs. forward scatter area (FSC-H/FSC-A) before identification of total T cells (B220$^-$CD3$^+$), granulocytes (B220$^-$CD11b$^+$Gr1$^+$), macrophages (B220$^-$CD11b$^+$F4/80$^+$), and B cells (F480$^-$B220$^+$). Cells were considered based on staining patterns: apoptotic cells were identified as Annexin V (Annexin V$^+$), necrotic cells as both Annexin V and 7-AAD (Annexin V$^+$7AAD$^+$), live cells as negative for both Annexin V and 7-AAD (Annexin V$^-$7AAD$^-$), and cell fragments or debris were characterized by high 7-AAD density.

Sup fig 6

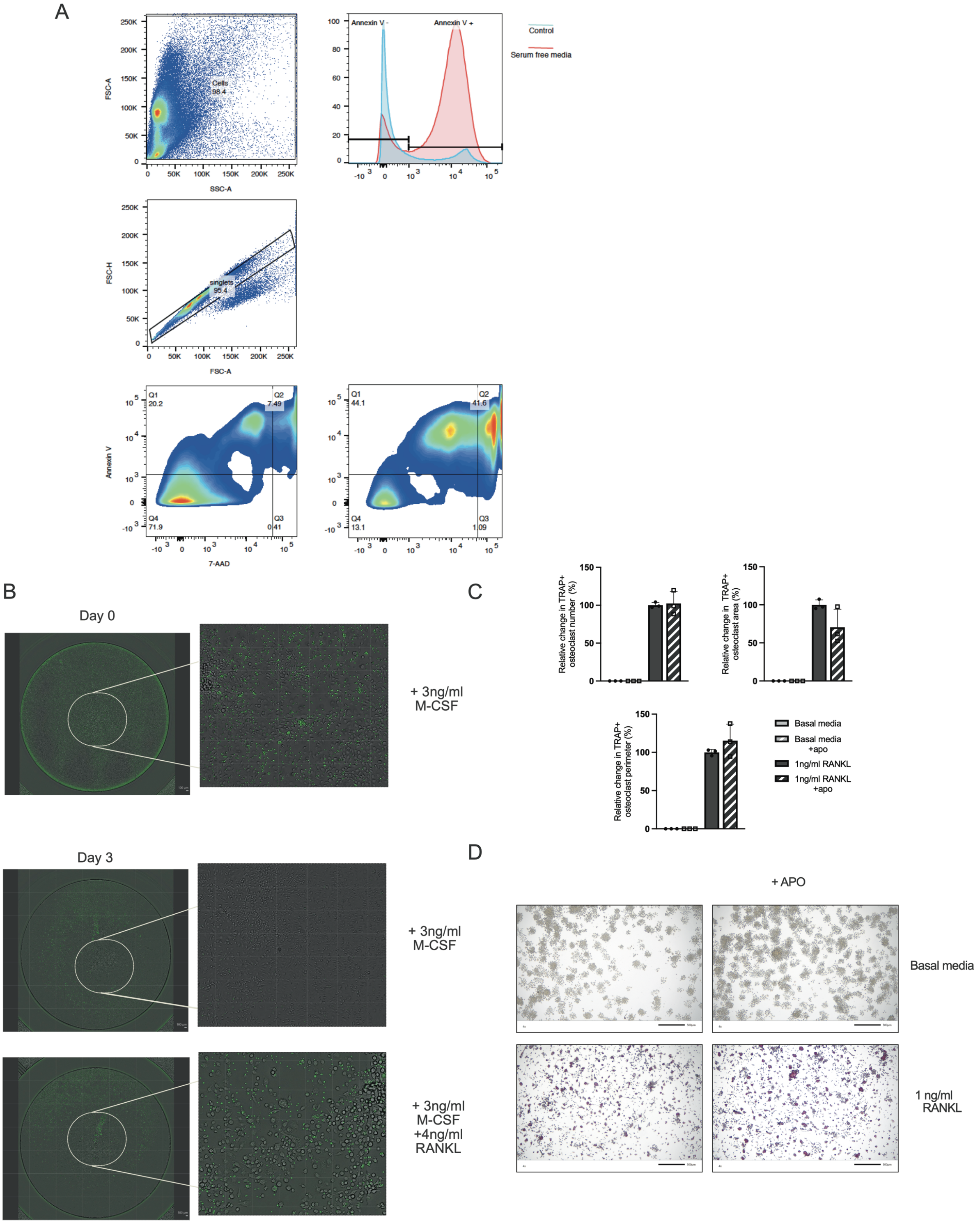


**Fig. S6: Phagocytosis assay.**

**A**. Representative flow cytometry gating strategy for identifying cell populations to validate thymocyte apoptotic ability for the phagocytosis assay. **B**. Validation of the long-term phagocytic capacity of osteoclasts using bone marrow macrophages as positive controls for internalizing FITC-conjugated apoptotic cells. **C**. Relative changes in the RAW 264.7 cells number, area, and perimeter of multinucleated TRAP+ cells after 3 days of differentiation, with or without apoptotic cells. **D**. Representative images of osteoclast differentiation after 3 days of stimulation with or without the presence of apoptotic cells. Sample sizes n=3.

| | Mean pixel density | | |
|---|---|---|---|
| **Serum marker** | **Control** | **2w** | **12w** |
| **Adiponectin/Acrp30** | 16751 | 25082 | 28711 |
| **BAFF/BLyS/TNFSF13B** | 10250 | 9321 | 29887 |
| **C-Reactive Protein/CRP** | 26830 | 31668 | 28801 |
| **CCL11/Eotaxin** | 8375 | 11935 | 8693 |
| **CCL21/6Ckine** | 22360 | 33290 | 35033 |
| **CD14** | 7935 | 24157 | 14496 |
| **Complement Component C5** | 8843 | 16344 | 13414 |
| **CX3CL1/Fractalkine** | 25061 | 36294 | 48671 |
| **CXCL16** | 20440 | 34220 | 40480 |
| **Cystatin C** | 21297 | 34783 | 7935 |
| **DPPIV/CD26** | 8182 | 12269 | 8419 |
| **Endostatin** | 31704 | 40265 | 41208 |
| **Flt-3 Ligand** | 8391 | 13124 | 12728 |
| **Gas 6** | 12630 | 22819 | 3569 |
| **IGFBP-5** | 9055 | 7939 | 7934 |
| **IL-6** | 7935 | 7935 | 8380 |
| **Leptin** | 8095 | 7892 | 7943 |
| **MMP-2** | 23824 | 28701 | 35774 |
| **MMP-3** | 20454 | 24611 | 29287 |
| **Osteoprotegerin** | 9013 | 11152 | 2280 |
| **Proliferin** | 7935 | 9892 | 9983 |
| **Resistin** | 19123 | 19190 | 19027 |
| **TNF-alpha** | 7935 | 7958 | 7952 |

Table S1 The mean pixel density of each serum marker

Changes in serum marker expression levels were determined based on the normalized pixel density.
